# Multitask Learning of Longitudinal Circulating Biomarkers and Clinical Outcomes: Identification of Optimal Machine-Learning and Deep-Learning Models

**DOI:** 10.1101/2023.08.19.553991

**Authors:** Min Yuan, Shixin Su, Haolun Ding, Yaning Yang, Manish Gupta, Xu Steven Xu

**Affiliations:** Department of Health Data Science, Anhui Medical University, Hefei, Anhui 230032, China; Department of Statistics and Finance, School of Management, University of Science and Technology of China; Clinical Pharmacology and Quantitative Science, Genmab Inc., Princeton, New Jersey, USA

**Author notes:** Corresponding author: Xu Steven Xu. MY and SS contributed equally.

**Keywords:** lung cancer, survival, machine learning, deep learning, longitudinal biomarkers

## Abstract

Many circulating biomarkers are assessed at different time intervals during clinical studies. Despite of the success of standard joint models in predicting clinical outcomes using low-dimensional longitudinal data (1-2 biomarkers), significant computational challenges are encountered when applying these techniques to high-dimensional biomarker datasets. Modern machine- or deep-learning models show potential for multiple biomarker processes, but systematic evaluations and applications to high-dimensional data in the clinical settings have yet to be reported. We aimed to enhance the scalability of joint modeling and provide guidance on optimal approaches for high-dimensional biomarker data and outcomes. We evaluated multiple deep-learning and machine-learning models using 24 clinical biomarkers and survival data from the SQUIRE trial, a phase 3 randomized clinical trial investigating necitumumab and standard gemcitabine/cisplatin treatment in patients with squamous non-small-cell lung cancer (NSCLC). Overall, we confirmed that longitudinal models enabled more accurate prediction of patients’ survival compared to those solely based on baseline information. Coupling multivariate functional principal component analysis (MFPCA) with Cox regression (MFPCA-Cox) provided the highest predictive discrimination and accuracy for the NSCLC patients with AUC values of 0.7 - >0.8 at various landmark time points and prediction timeframes, outperforming recent advanced Transformer and convolutional neural network deep-learning algorithms (TransformerJM and Match-Net, respectively). In conclusion, we identified that MFPCA-Cox represents a robust and versatile joint modeling algorithm for high-dimensional biomarker longitudinal data with irregular and missing data, capturing complex relationships within the data, yielding accurate predictions for both longitudinal biomarkers and survival outcomes, and gaining insights into the underlying dynamics.

## Introduction

The essential elements of precision medicine lie in the accurate and precise prediction of treatment response, toxicity, early disease progression, as well as good understanding of mechanism of actions, so that personalized medical and treatment decisions can be available for individual patients [1, 2]. Achieving this goal necessitates harnessing all available data. Unfortunately, most models in clinical studies fail to fully utilize longitudinal data and instead rely solely on a snapshot of baseline information.

Collecting circulating laboratory and biological markers in clinical trials is frequently done in oncology research. A large number of circulating markers are assessed at different intervals during the trial, such as immune profiling markers, cytokines, blood counts, renal and hepatic function tests, coagulation and lipid panel, growth factor proteins, inflammation markers, and others, often in conjunction with imaging and other clinical evaluations. Pharmacokinetic concentrations are also typically measured at multiple time points. Furthermore, circulating tumor DNA (ctDNA) is an emerging marker to identify presence of cancer and monitor treatment response. Previous analyses have shown the significant contributions of longitudinal information to risk estimation [3-7].

The current analysis framework for on-treatment longitudinal trajectories biomarkers and survival outcomes is often limited to a specific biomarker or a small set of biomarkers chosen based on prior knowledge or hypothesis, which may overlook important changes in other biomarkers that are relevant for predicting treatment response or toxicity. To gain a comprehensive understanding of disease mechanisms, treatment response, disease progression, and adverse event dynamics, it is necessary to integrate longitudinal data of all biomarkers collected during clinical studies. However, due to the complexity of on-treatment biomarker trajectories, there are limited tools available to model the trajectories of a large number of biomarkers and clinical outcomes simultaneously. This requires more advanced modeling approaches to capture complex relationships among different biomarkers over time and between biomarker changes and clinical outcomes accurately.

In recent years, there has been an explosion of machine and deep learning approaches, with many of them incorporating longitudinal or time-series data to predict outcomes [8-13]. Nevertheless, no studies have assessed the performance of these methods concerning high-dimensional longitudinal biomarker processes and clinical outcomes. Therefore, further exploration and evaluation are warranted in this context. The objective of this study was to address the limitations of current biomarker modeling and learning frameworks and, for the first time, to identify optimal machine- and/or deep-learning algorithms that can enable multitask learning and simultaneous modeling of high-dimensional longitudinal biomarker trajectories over time and clinical outcomes together.

## Materials and Methods

### Patient Data Set

The analysis in this study utilized data from the SQUIRE clinical trial, which was a phase 3, multicentered, open-label randomized clinical trial [14]. The main objective of the trial was to investigate whether adding necitumumab to the standard gemcitabine and cisplatin treatment would result in improved outcomes for patients with stage IV squamous non-small-cell lung cancer.

The trial compared two treatment groups: the experimental arm, which received a combination of necitumumab, gemcitabine, and cisplatin, and the control arm, which received only gemcitabine and cisplatin as the first-line therapy.

For this particular analysis, data from the control arm were used. A total of 548 patients diagnosed with stage IV squamous-cell NSCLC, who had not previously received treatment for metastatic disease, were randomly assigned to the control arm. The chemotherapy regimen involved administering gemcitabine at a dose of 1250 mg/m²intravenously over 30 minutes on days 1 and 8, along with cisplatin at a dose of 75 mg/m²intravenously over 120 minutes on day 1. The treatment was given in cycles of 21 days, and a maximum of six cycles were administered.

Longitudinal measurements of 24 circulating biomarkers, including, hematology, coagulation, and serum chemistry profiles, were collected from laboratory tests (Supplementary Table S1). Hematology and serum chemistry profiles were assessed at various time points (Supplementary Figure S1), including screening before randomization, the start of week 1, and week 2 of every 3-week cycle, at the end of therapy, and during a 30-day follow-up period. The coagulation profile was evaluated at screening before randomization, the start of week 1, week 4, at the end of therapy, and during the 30-day follow-up. If clinically indicated, the coagulation profile was assessed more frequently, and any unscheduled measurements were assigned to the nearest treatment cycle. One patient with missing circulating biomarker measurements was excluded from the analysis, leaving a total of 547 patients included for the study.

The primary efficacy endpoint in this study was overall survival time (OS), which refers to the duration from randomization to the date of death from any cause. Patients who were still alive at the conclusion of the study or lost to follow-up were considered censored data in the analysis.

The clinical trial protocol obtained ethical approval from the review boards at each of the participating sites, adhering strictly to the principals outlined in the Declaration of Helsinki and Good Clinical Practice guidelines. Before enrolling in the study, all patients provided written informed consent, demonstrating their voluntary and well-informed decision to participate in the research. The trial was registered on www.ClinicalTrials.gov under the unique identifier NCT00981058.

### Predictive models for survival using baseline biomarkers

We first evaluated baseline biomarker models to serve as benchmarks for comparison with longitudinal models. Traditional Cox proportional hazards model and 6 different deep-learning survival models, i.e., Logistic-Hazard [15], Cox-Time and Cox-CC [16], DeepHit [17], PC Hazard [18], and DeepSurv [19], were utilized to evaluate the association between baseline biomarkers and NSCLC patients’ survival. Details of the deep survival neural networks are provided in supplementary. Alternatively, we also performed dimensionality reduction of the baseline measurements of the 24 circulating biomarkers using principal component analysis (PCA). Subsequently, the top principal components that collectively accounted for over 99% of the overall variance were included as the explanatory variables in Cox model and the deep-learning survival models.

### Multitask learning of biomarker trajectories and survival

The workflow for data processing, deep- and machine-learning based multitask leaning, predictive performance evaluation, and dynamic prediction is illustrated in Figure 1. The primary goal of this study is to simultaneously learn from longitudinal trajectories of multiple biomarkers and dynamically predict the biomarkers longitudinal profiles and survival of patients with advanced non-small cell lung cancer. To achieve this, we employ both machine learning and deep learning techniques, as described as follows:

**Figure 1.**
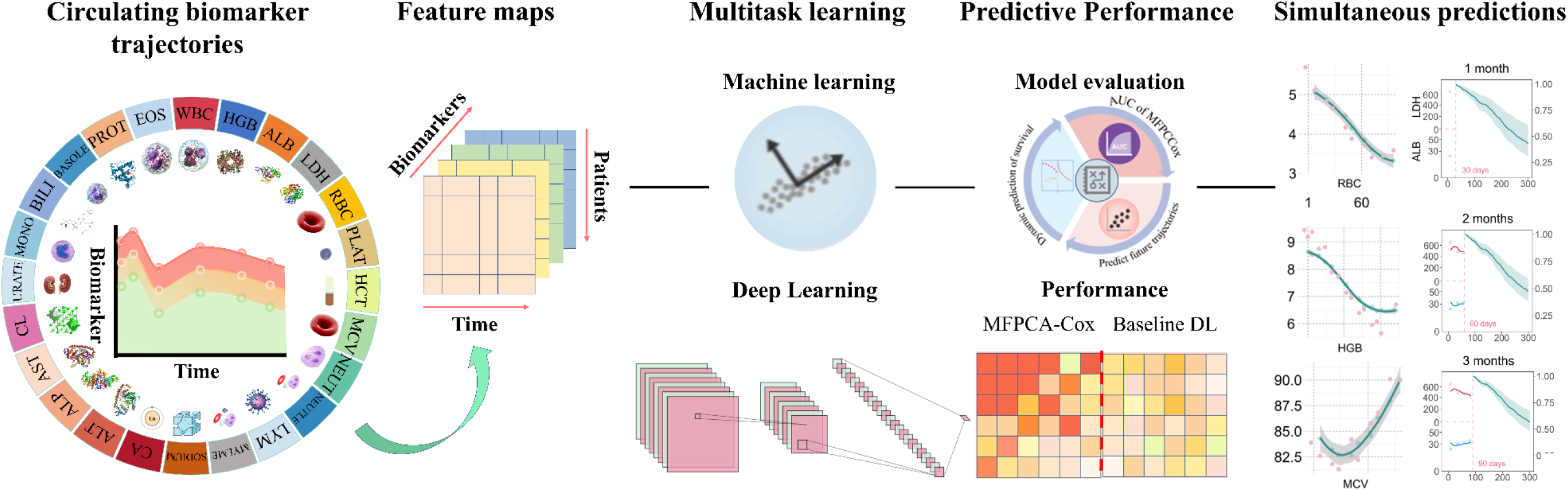
Workflow of Multitask Learning of Longitudinal Circulating Biomarkers and Clinical Outcomes using Machine-Learning and Deep-Learning Models.

#### Match-Net

Match-Net is a Convolutional Neural Network (CNN) specially designed to incorporate multiple longitudinal variables for precise survival prediction [20]. Match-Net directly estimates the failure probability or the likelihood of an event occurring within predefined prediction windows. The final output of Match-Net is a vector, with each element corresponding to one of the contiguous prediction windows of interest. To ensure interpretability, a softmax function is applied to the vector, ensuring that the failure probabilities across all prediction windows sum up to one.

#### TransformerJM

TransformerJM is a more recent neural network designed for predicting longitudinal and survival trajectories, akin to natural language processing tasks [21]. It transforms sequences of longitudinal measurements into future longitudinal and survival predictions using a sequence-to-sequence model. The model’s key feature is the attention mechanism, which identifies relevant context between sequence elements, allowing it to consider dependencies between visits for a comprehensive patient profile. Multi-head attention is used to calculate attention in parallel, enabling more refined representations of a patient’s disease status. Positional encodings are added to give the model a sense of order in longitudinal data by representing visit times. The architecture comprises “decoder” blocks, each consisting of masked multi-head self-attention followed by a feed-forward neural network. This design enables the model to learn interactions between longitudinal outcomes and incorporate contextual information from previous visits. For predicting future visits, the model adds positional encoding of the desired prediction time to the last visit’s vector and passes it through a final decoder block to generate a single vector predicting the patient’s state at that time. Training involves minimizing two loss functions: longitudinal loss (mean squared error) and survival loss (negative log likelihood function). This powerful architecture leverages attention to capture dependencies between visits and provides accurate longitudinal and survival predictions for patient outcomes.

#### MFPCA-Cox

MFPCA-Cox is a two-stage machine learning method that performs feature extraction based on multivariate functional principal component analysis (MFPCA) [22]. In the first stage, the clinical biomarker trajectories are considered as stochastic functions over the longitudinal visit times. MFPCA is used to reduce the dimensionality of the longitudinal biomarker data while capturing the essential patterns of variability among the biomarkers by extracting the changing patterns (features) of multiple biomarker trajectories. Karhunen–Loève decomposition was used to project the trajectory profiles of individual biomarkers onto a set of basis functions, resulting in a set of functional principal component scores (FPCs) for each biomarker [23]. Top FPCs that collectively captured 99% of the variability, serving as essential features of longitudinal biomarkers, were retained for further analysis. In the second stage, these features were used to extrapolate biomarker trajectories and scores on these features were used as predictors in a Cox proportional hazards model to model the time-to-event data, such as survival outcomes and conduct predictions of outcomes over time. This personalized dynamic prediction framework can be updated as new observations collected to reflect the patient’s latest prognosis.

#### MFPCA-DL Survival models

Similarly, we evaluated the combinations of MFPCA with the 6 different deep-learning survival models, i.e., Logistic-Hazard [15], Cox-Time [16], DeepHit [17], Cox-CC [16], PCHazard [18], and DeepSurv [19]. A brief introduction to these methods is included in the supplementary material.

### Model training and evaluation

To assess model performance, we employed 5-fold cross-validation. For survival outcome, the predictive ability of the models was evaluated using the Brier score [24] and the area under the receiver operating characteristic curve (AUC). The Brier score measures the accuracy of predicted survival probabilities, while the AUC measures the models’ ability to distinguish between patients who will survive or perish within a specific timeframe. For longitudinal data for biomarkers, consistency between predicted and observed data was assessed.

### Dynamic Prediction

The landmark models’ predictive capabilities were demonstrated using patient cases with varying biomarker trajectories. Conditional probabilities, denoted as *p*(*t* | *s*), were predicted to estimate a patient’s likelihood of survival or death at a future time t, based on biomarker values measured up to time s (s = 1, 2, 3, 4, 5 and 6 months). To achieve reliable estimates, 1000 random samples from conditional distribution *p*(*t* | *s*) were simulated, resulting in 95% Prediction Intervals (PIs) for each prediction. These simulations effectively showcased how a patient’s risk of death changes over time in relation to their marker values.

## Results

### Patient characteristics

Baseline characteristics of demographic variables of 547 eligible patients treated with gemcitabine and cisplatin were summarized in Supplementary Table S2. Briefly, most of the patients were smokers (494 [90.3%]), males (457 [83.5%]) and white (455 [83.2%]); 474 [86.7%] patients were recruited from North America, Europe and Australia; 180 [32.9%] patients had an ECOG performance score of 0 and 320 [58.5%] patients had an ECOG performance score of one. The median duration of follow-up and median survival were 4.23 months (IQR 2.35–4.92) and 9.20 months (IQR 5.24–16.62) respectively. In total, 442 patients had died with a censoring rate 80.8% at the end of the follow-up. The Kaplan-Meier plots of the training and testing datasets from a randomly selected cross-validation fold are illustrated in Supplementary Figure S2.

### Survival information embedded in longitudinal biomarker data

Figure 2 presents the trajectories of 24 circulating biomarkers for both surviving and deceased patients. Clear distinctions in the trajectories between the two groups were evident. Notably, the WBC level exhibited higher volatility in the deceased group, while the ALB consistently showed higher levels in the survival group compared to the deceased group. Interestingly, the LDH index displayed an opposite trend, with the deceased group consistently exhibiting higher LDH measurements compared to the survival group. These observations highlight the diverse patterns of biomarkers in relation to different survival outcomes.

**Figure 2.**
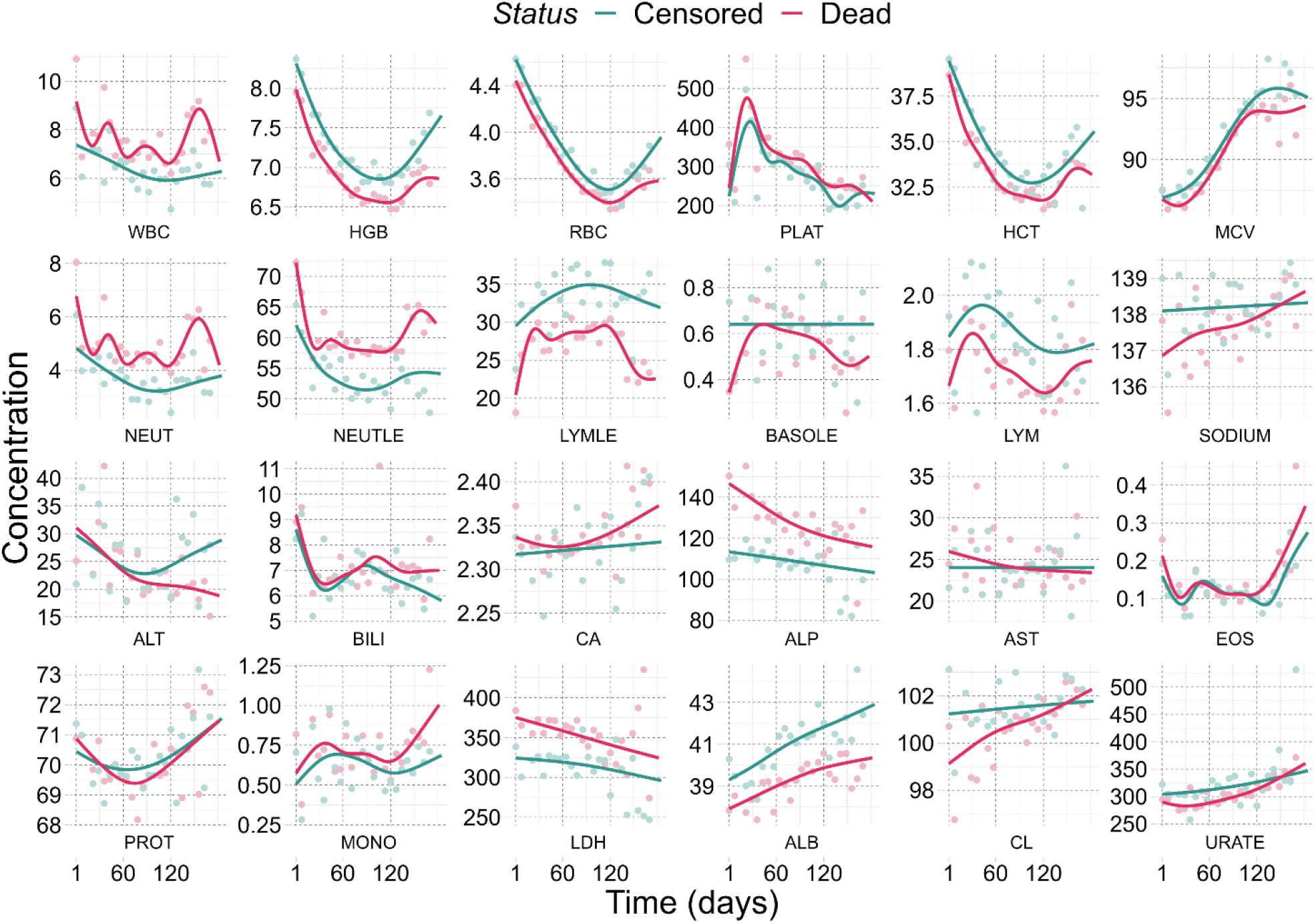
Different evolution of longitudinal biomarker profiles over time between death vs. alive.

### Predictive performance of baseline biomarkers for survival

We investigated and compared the associations between baseline circulating biomarker measurements and survival outcomes using both Cox regression and various deep learning models. To visualize the performance of these baseline models, a heat-map was utilized, displaying their Area under the Curve (AUC) and Brier Score (BS) at different landmark time points (see Supplementary Figure S3).

In general, the baseline deep-learning models demonstrated certain predictive capabilities for overall survival, with AUCs ranging from 0.60 to 0.70 (Supplementary Figure S3). Among the six deep-learning models, DeepSurv appeared to provide the highest AUC (discrimination capability) and BS (prediction accuracy). However, the traditional Cox model with baseline circulating biomarker measurements exhibited relatively poor performance, with AUCs ranging from 0.5 to 0.6. Consistently, the Cox model exhibited lower prediction accuracy (measured by BS) compared to the deep-learning models. However, upon further investigation, we discovered significant collinearity among the baseline biomarker measurements, which seemed to be the reason behind the suboptimal performance of the Cox model (Supplementary Figure S4). To address this issue, we employed PCA for dimension reduction, aiming to alleviate the collinearity. Subsequently, we used the PCA-extracted features to construct both the Cox and deep-learning models. After applying dimension reduction, the performance of the Cox model improved significantly, reaching a level comparable to that of the deep-learning models (Supplementary Figure S5). Among the deep-learning survival models, DeepSurv remained the top performance. This experiment confirmed our hypothesis that traditional Cox models are less robust against collinearity in the data compared to deep-learning approaches.

### Predictive performance of biomarker trajectories for survival

To assess the predictive performance of on-treatment longitudinal biomarkers for clinical outcomes, we employed landmark modeling at different landmark time points, using both deep-learning and machine-learning approaches. The results clearly indicate that incorporating on-treatment biomarker trajectory measurements significantly enhanced predictive performance for survival, as evident from improved AUC and BS compared to baseline Cox and DeepSurv models, which solely relied on baseline biomarker measurements (Figure 3).

**Figure 3.**
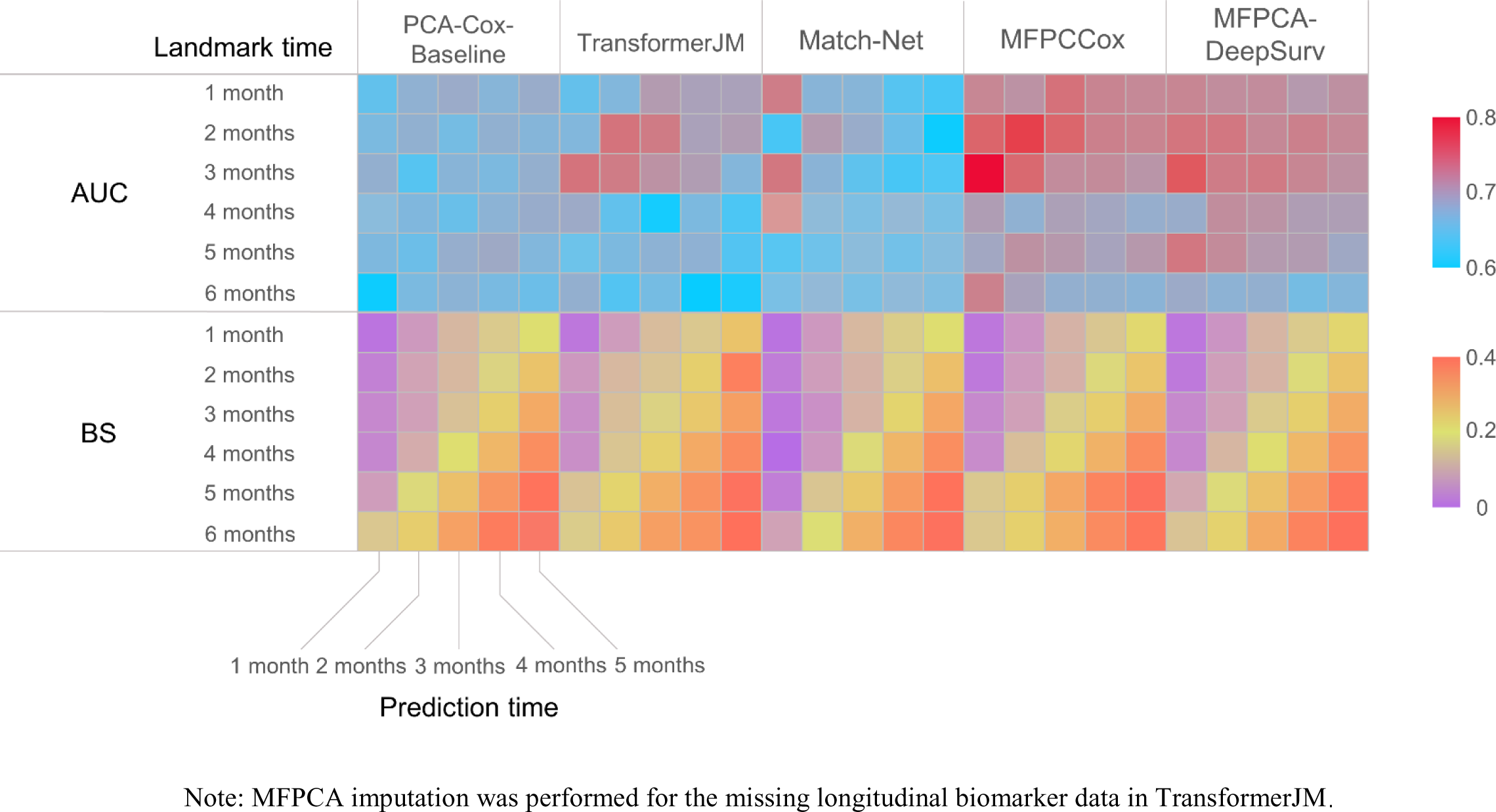
Predictive performance of deep-learning and machine-learning approaches for clinical outcome.

All machine- and deep-learning methods utilizing longitudinal data demonstrated substantial improvements over the baseline models. However, notably, while TransformerJM and Match-Net are popular deep-learning methods, their performances showed only modest improvements over the baseline PCA-Cox model, exhibiting better AUC and BS at early landmark time points (1-3 months). Interestingly, the MFPCA-Cox and MFPCA-DeepSurv models achieved the highest performance, exhibiting AUC values greater than or close to 0.8. This finding underscores the effectiveness of incorporating MFPCA for feature extraction, as it significantly enhances the predictive accuracy of the models.

### Prediction of biomarker trajectories using MFPCA

The MFPCA model not only excels at extracting meaningful features from longitudinal data encompassing multiple circulating biomarkers but also proficiently learns the longitudinal patterns of these biomarkers, offering predictions for their trajectory measurements. To assess the predictive capabilities of MFPCA in generating biomarker trajectories, we conducted a comparative analysis between the predicted biomarker measurements by MFPCA and the actual measurements, represented through scatter plots for all 24 circulating biomarkers (Figure 4).

**Figure 4.**
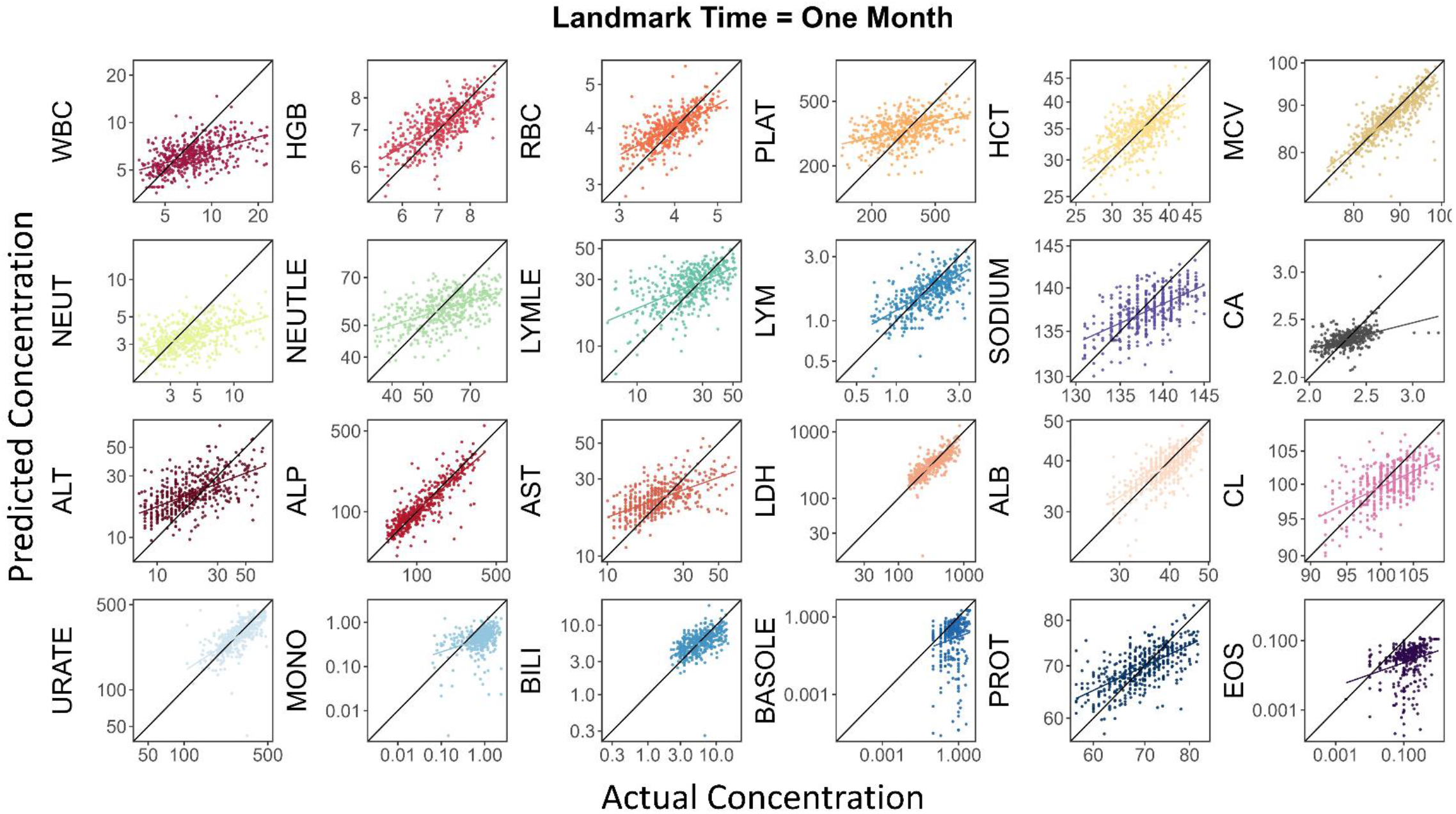
MFPCA adequately predict multiple biomarker trajectories at the same time.

The comparison revealed that MFPCA demonstrates impressive accuracy in predicting the trajectories of most biomarkers, including HGB, RBC, HCT, MCV, LYM, ALP, LDH, ALB, URATE and BILI, etc. The scatter points closely cluster around the diagonal, indicating a favorable alignment between the predicted and actual trajectories for these biomarkers. These results suggest that MFPCA effectively captures the inherent patterns and trends of the biomarkers, rendering it a dependable method for trajectory prediction.

Moreover, to investigate MFPCA’s performance at an individual level, we randomly selected a patient and examined the trajectories predicted by MFPCA. Remarkably, MFPCA successfully reconstructed the biomarker trajectories with high precision, as shown in Figure 5. This observation further emphasizes the MFPCA model’s ability to generate accurate trajectory predictions for a large number of biomarkers for individual patients at once.

**Figure 5.**
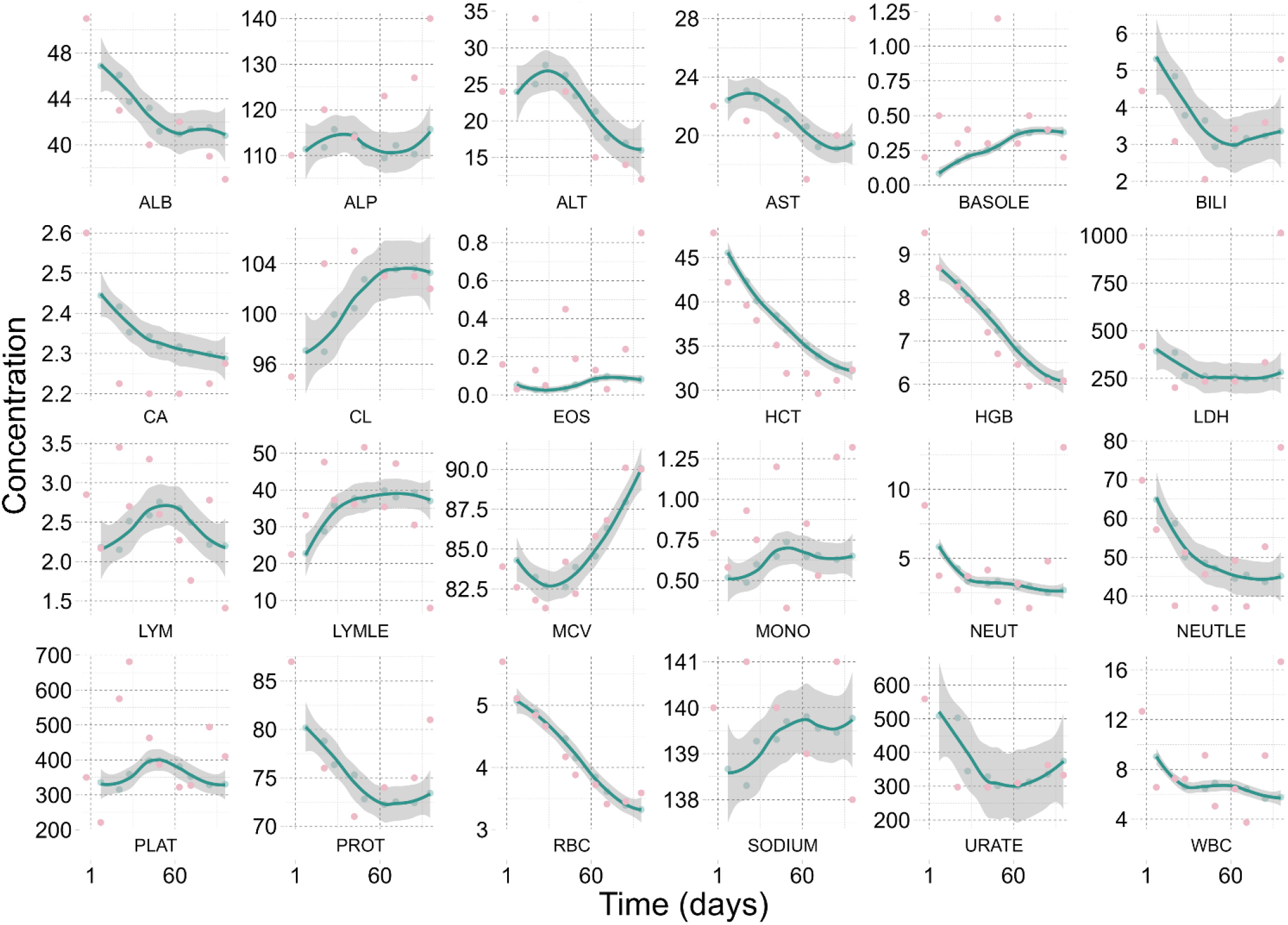
Prediction of biomarker trajectories for an individual patient.

### Interpretability of MFPCA

To further investigate the interrelationships among FPCA features, we employed network analysis and visualized the connections among the functional principal components (FPCs) from the longitudinal biomarker trajectories (Figure 6). By aggregating the FPCs that collectively explained 99% of the variation for the longitudinal data of each biomarker, we computed pairwise correlation coefficients. Using a threshold of 0.5, we constructed a network diagram (an adjacency matrix) to depict the relationships among the FPCs. This visualization revealed significant correlations among FPCs of various biomarkers at different landmark time points. As expected, the first principal component of sodium exhibited a positive correlation with the first principal component of chloride, and the first two principal components of AST and ALT were positively correlated as well. Additionally, WBC and NEU showed a positive correlation.

**Figure 6.**
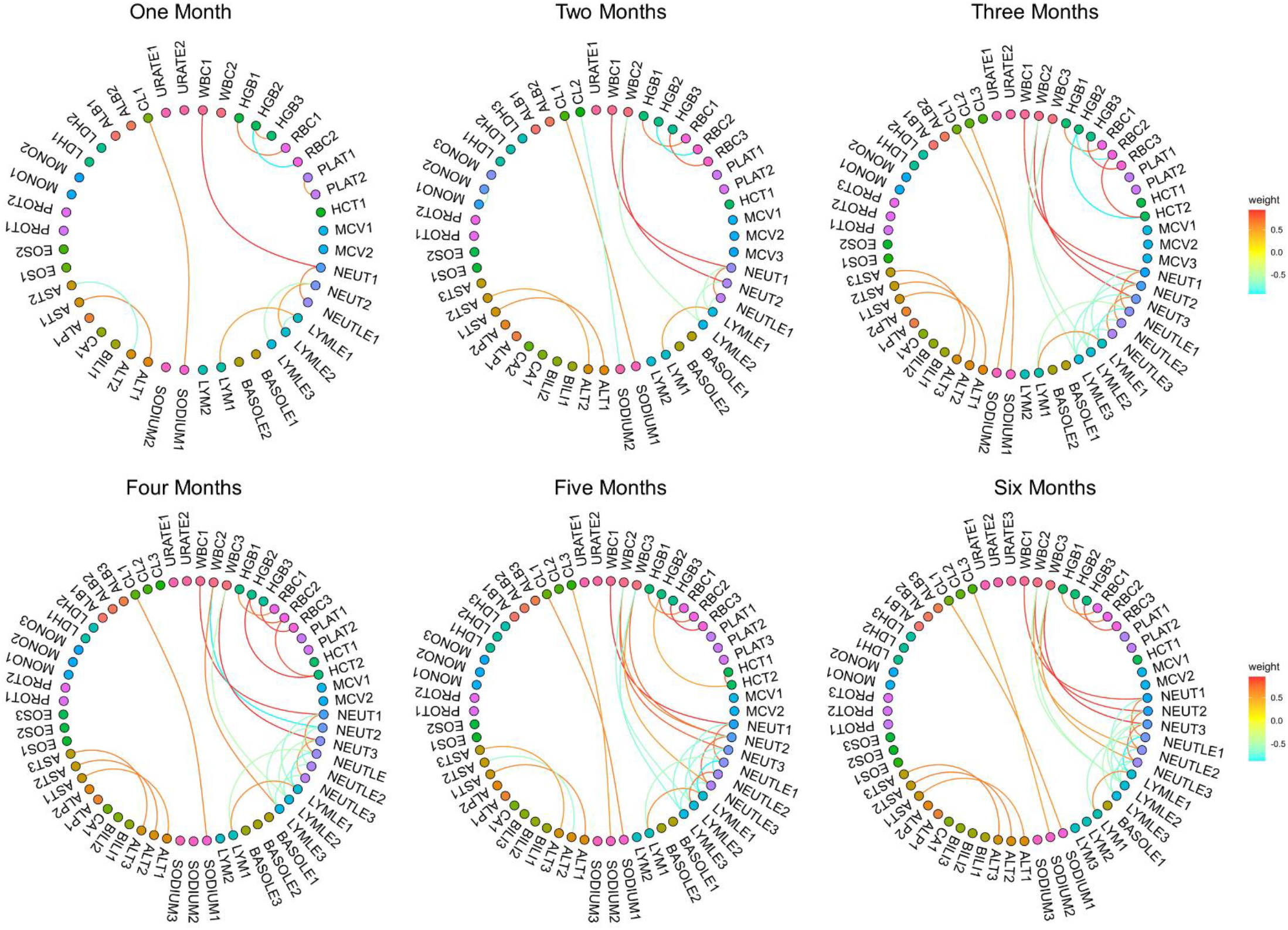
Network analysis to depict relationships among FPCA features.

Subsequently, we conducted a multivariate survival analysis at each landmark to identify the most relevant features from the multiple circulating biomarkers. The p-values were ranked in ascending order for comparison, and the results were presented in Figure 7. LDH, ALB, NEUT/NEUTLE/WBC, HGB/RBC, MCV, and ALT are repeatedly identified as significant biomarkers in the landmark models.

**Figure 7.**
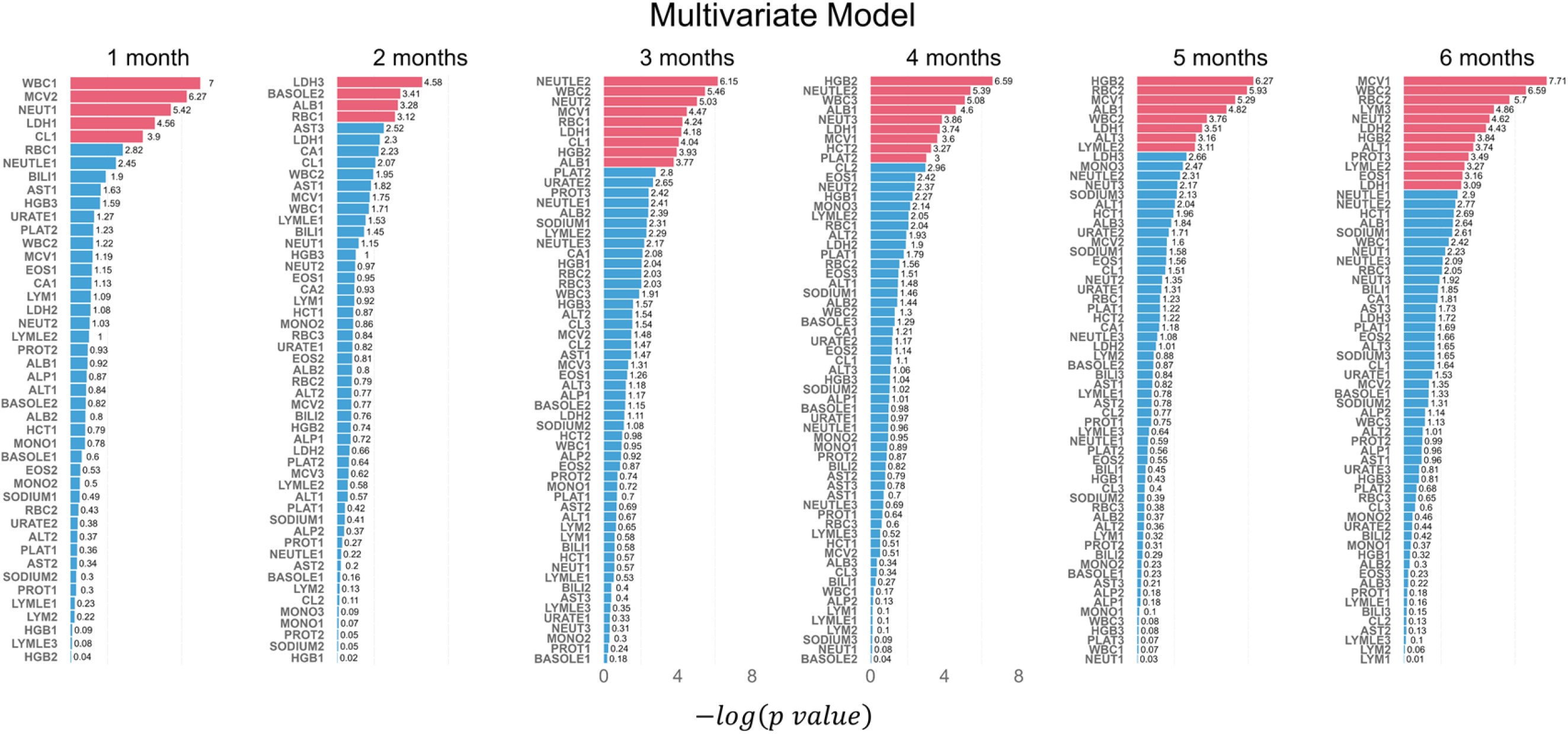
Network analysis to depict relationships among FPCA features.

### Dynamic prediction of survival using longitudinal biomarker data

Figure 8 shows the dynamic predictions of survival for two representative patients. To provide a clear illustration, we display the trajectories of LDH and ALB, which are the two most frequently identified significant longitudinal biomarkers, highlighting their joint dynamic predictions for survival.

**Figure 8.**
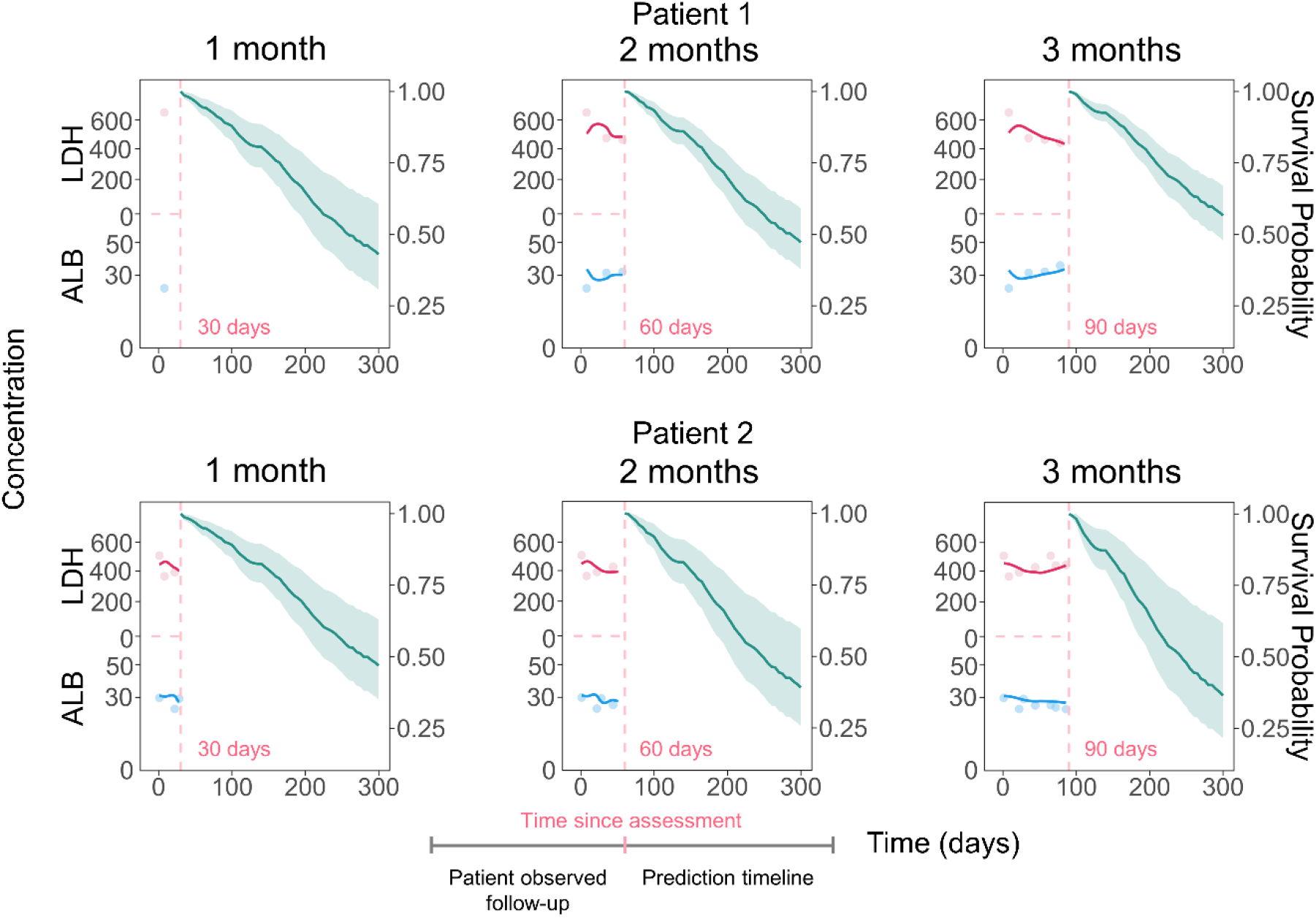
Simultaneous dynamic prediction of biomarkers and survival outcome.

The first patient’s LDH trajectory demonstrates a decreasing trend over time, while the ALB trajectory exhibits an increasing trend. Conversely, the second patient’s LDH levels show an elevation, while their ALB levels display a declining pattern across time. Notably, the dynamic predictions for the first patient suggest an improvement in survival as time progresses, while the predictions for the second patient indicate a decline in survival. These dynamic trends of biomarkers offer valuable insights into the patients’ prognoses, based on the changing patterns of LDH and ALB over time.

## Discussion

Longitudinal circulating biomarker measurements are essential in clinical studies for multiple reasons. They enable the assessment of treatment efficacy, prediction of treatment response, and early detection of disease progression. Additionally, they aid in monitoring drug toxicity and contribute to personalized medicine, tailoring treatments to individual patients. The analysis of circulating biomarkers holds significant potential in improving cancer diagnosis, treatment, and patient outcomes, enhancing cancer care through better understanding of disease biology and more effective, patient-centric medical interventions. As anticipated, our research has revealed that forecasting survival in patients with metastatic NSCLC can be significantly improved by analyzing additional longitudinal biomarker information, rather than relying solely on baseline variables. The longitudinal approach enables a dynamic and more accurate determination of patient’s risk.

Numerous studies have confirmed the correlation between longitudinal biomarkers and various outcomes [25-28]. Currently, the joint models typically utilize individual models for each data trajectory, such as linear mixed models for longitudinal data and the Cox proportional hazard model for survival analysis [29]. However, despite their effectiveness, the standard joint models encounter significant computational challenges when applied to large datasets, especially when the dimensionality of the random effects component increases [29]. These computational limitations can hinder their applicability to extensive datasets, emphasizing the need for more efficient approaches or alternative methods to handle the complexities of high-dimensional data in joint modeling scenarios. Our study aimed to significantly enhance the scalability of joint modeling of longitudinal biomarkers and clinical outcomes and provide guidance on machine- and deep-learning approaches that are best suited for predictions of high-dimensional biomarker data and outcomes.

We evaluated multiple deep learning and machine learning based multitask learning frameworks for longitudinal biomarker trajectories and clinical outcomes. To our knowledge, this is the first study attempting to identify optimal machine- and deep-learning algorithms for high-dimensional biomarker and clinical outcome data collected in clinical studies. Our study demonstrated that MFPCA could effectively capture relationships between a large number of biomarker trajectories by learning representations and extracting informative features from the complex longitudinal data. Coupling with Cox modeling, the MFPCA-Cox workflow can generate predictive distributions for both longitudinal and time-to-event data, enabling the simultaneous forecasting of multiple biomarker trajectories of interest, and informing the expected survival time. Our analysis on the actual clinical data showed that MFPCA-Cox provided the highest predictive discrimination and accuracy for NSCLC patients with AUC values of 0.7 - >0.8 at various landmark time points and prediction timeframes.

In clinical trials, patients are monitored over a specific period (e.g., weeks to months). While biomarker measurements are scheduled at specific fixed time points, the collection of samples often occur at irregular intervals due to delayed or missing visits, resulting in a notable amount of missing data. The deep-learning networks require observations to lie on a grid of fixed time steps [30], so the observed data is first rounded to the nearest time corresponding to an established grid. In addition, missing values are not allowed in the deep-learning networks. Imputations based on the last observed measurement or other sophisticated models are required. On the other hand, MFPCA does not have restrictions on fix time intervals and allows missing longitudinal data. In fact, FPCA is often used to impute missing data in longitudinal studies [31, 32]. In our TransformerJM modeling, we actually employed MFPCA to impute missing data points and to create modified input data into the TransformerJM network. Our analysis indicated that FPCA-imputation of the longitudinal biomarker data improved the predictive performance of TransformerJM (Supplementary Figure S6).

While survival prediction, or prognosis, has traditionally been the primary focus in medical research [33, 34], there is a growing emphasis in precision medicine on forecasting disease-or treatment-related biomarker trajectories. This involves utilizing temporal correlations and associations between biomarkers to predict their evolution over time. Modeling on-treatment biomarker trajectories, particularly high-dimensional biomarkers simultaneously can be challenging, given different distributions and evolutions of different biomarkers. Currently, the analysis of high-dimensional on-treatment biomarker trajectories lacks standardized methods. MFPCA demonstrated impressive versatility and effectively handled all the dynamic biomarkers data from the NSCLC dataset, capturing intricate relationships and time-varying patterns inherent in the data and providing adequate predictions for most biomarkers.

In conclusion, MFPCA-Cox represents a robust and versatile joint modeling algorithm for high-dimensional biomarker longitudinal data with irregular and missing data, capturing complex relationships within the data, yielding accurate predictions for both longitudinal biomarkers and survival outcomes, gaining insights into the underlying dynamics, and enabling better clinical decision-making and patient care.

## ACKNOWLEDGMENT

This work was partially supported by the Natural Science Foundation of Anhui Province (No. 2008085MA09), the National Natural Science Foundation of China (No. 11671375).

## Supplementary Materials

### Part 1. Deep-learning survival Models

#### DeepSurv

DeepSurv [19] is a deep learning model designed for survival analysis that extends the Cox proportional hazards model using neural networks. With its multi-layered architecture and non-linear activation functions, DeepSurv captures complex relationships between covariates and survival times. It effectively handles censored data, making it applicable in real-world scenarios where complete information is unavailable. The model is trained end-to-end with a custom loss function, ensuring optimal performance and avoiding overfitting. DeepSurv provides a flexible and powerful tool for analyzing high-dimensional survival data. Its versatility and capability to handle complex scenarios make it valuable for a wide range of research and clinical applications.

#### Logistic-Hazard Model

The Logistic-Hazard model [18] is a hybrid approach that combines logistic regression with survival analysis techniques. It aims to predict binary outcomes (e.g., survival vs. non-survival) while considering the time-to-event information. This model is suitable for scenarios where the event of interest occurs over time, and there is interest in understanding the factors influencing both the binary outcome and the time it takes to reach the event.

#### Cox-Time Model

The Cox-Time model [16] is an extension of the traditional Cox proportional hazards model used for survival analysis. It addresses time-varying covariates by allowing the coefficients to vary over time, thus capturing the dynamic nature of the predictors’ impact on survival. The Cox-Time model is valuable when the relationship between covariates and survival changes throughout the observation period.

#### DeepHit

DeepHit [17] is a deep learning model specifically designed for survival analysis tasks. It combines a recurrent neural network (RNN) with a multi-layer perceptron (MLP) to handle censored survival data effectively. DeepHit can predict the entire survival curve, providing a more comprehensive view of the event probabilities over time, making it suitable for complex survival scenarios.

#### Cox-CC Model

The Cox-CC (Concordance-Counting) model [16] is an extension of the traditional Cox proportional hazards model that incorporates additional information from the counting process framework. By considering the concordance between predicted and observed event times, the Cox-CC model aims to improve prediction accuracy for time-to-event outcomes.

#### Piecewise Constant Hazard (PCHazard) Model

The Piecewise Constant Hazard (PC-Hazard) model [18] is a survival analysis method that approximates the hazard function by assuming it remains constant within predefined time intervals. The model breaks the observation period into segments and estimates a constant hazard for each segment. This approach allows for a more flexible representation of the hazard function compared to traditional parametric models, capturing abrupt changes in risk over time. Researchers commonly use the PC-Hazard model when they suspect that the hazard function varies in distinct intervals. It is applicable in various fields, including medical research, engineering, and social sciences, where survival or time-to-event data are analyzed. However, as with any statistical model, careful interpretation and consideration of the model’s assumptions are crucial for accurate results. The choice of the number and locations of breakpoints can significantly impact the model’s performance and should be based on the specific data characteristics and domain knowledge.

### Part 2. Supplementary Tables and Figures

**Table S1:** Circulating biomarkers and their abbreviation.

| Abbreviation | Biomarker | PARCAT |
| --- | --- | --- |
| ALB | Albumin | Serum Chemistry |
| ALP | Alkaline Phosphatase | Serum Chemistry |
| ALT | Alanine Aminotransferase | Serum Chemistry |
| AST | Aspartate Aminotransferase | Serum Chemistry |
| BASOLE | Basophils/Leukocytes | Hematology |
| BILI | Bilirubin | Serum Chemistry |
| CA | Calcium | Serum Chemistry |
| CL | Chloride | Serum Chemistry |
| EOS | Eosinophils | Hematology |
| HCT | Hematocrit | Hematology |
| HGB | Hemoglobin | Hematology |
| LDH | Lactate Dehydrogenase | Serum Chemistry |
| LYM | Lymphocytes | Hematology |
| LYMLE | Lymphocytes/Leukocytes | Hematology |
| MCV | Ery. Mean Corpuscular Volume | Hematology |
| MONO | Monocytes | Hematology |
| NEUT | Neutrophils | Hematology |
| NEUTLE | Neutrophils/Leukocytes | Hematology |
| PLAT | Platelet | Hematology |
| PROT | Protein | Serum Chemistry |
| RBC | Erythrocytes | Hematology |
| SODIUM | Sodium | Serum Chemistry |
| URATE | Urate | Serum Chemistry |
| WBC | Leukocytes | Hematology |

**Table S2.** Baseline characteristics for the study participants.

| **Category/statistic** | **Gemcitabine + Cisplatin (N = 547)** |
| --- | --- |
| **Age at study** |  |
| n | 547 |
| >= 18 - < 65 years | 339 (62.0%) |
| >=65 - < 70 years | 111 (20.3%) |
| >=70 years | 97 (17.7%) |
| **SEX** |  |
| n | 547 |
| Male | 457 (83.5%) |
| Female | 90 (16.5%) |
| **RACE** |  |
| n | 547 |
| White | 455 (83.2%) |
| Non-White | 92 (16.8%) |
| **Geographical region** |  |
| n | 547 |
| North America, Europe, and Australia | 474 (86.7%) |
| South America, South Africa, and India | 55 (10.0%) |
| Eastern Asia | 18 (3.3%) |
| **Smoking history** |  |
| n | 547 |
| Smoker | 494 (90.3%) |
| Ex-Light Smoker | 26 (4.9%) |
| Non Smoker | 27 (4.8%) |
| **ECOG Performance Status** |  |
| n | 547 |
| 0 | 180 (32.9%) |
| 1 | 320 (58.5%) |
| 2 | 47 (8.6%) |
| **Baseline BMI (kg/m<sup>2</sup>)** |  |
| n | 547 |
| MEAN (SD) | 24.753 (4.854) |
| MEDIAN (min, max) | 24.435 (14.872, 55.820) |
| **Baseline BSA (m<sup>2</sup>)** |  |
| n | 547 |
| MEAN (SD) | 1.803 (0.206) |
| MEDIAN (min, max) | 1.800 (1.3, 2.52) |

**Figure S1:**
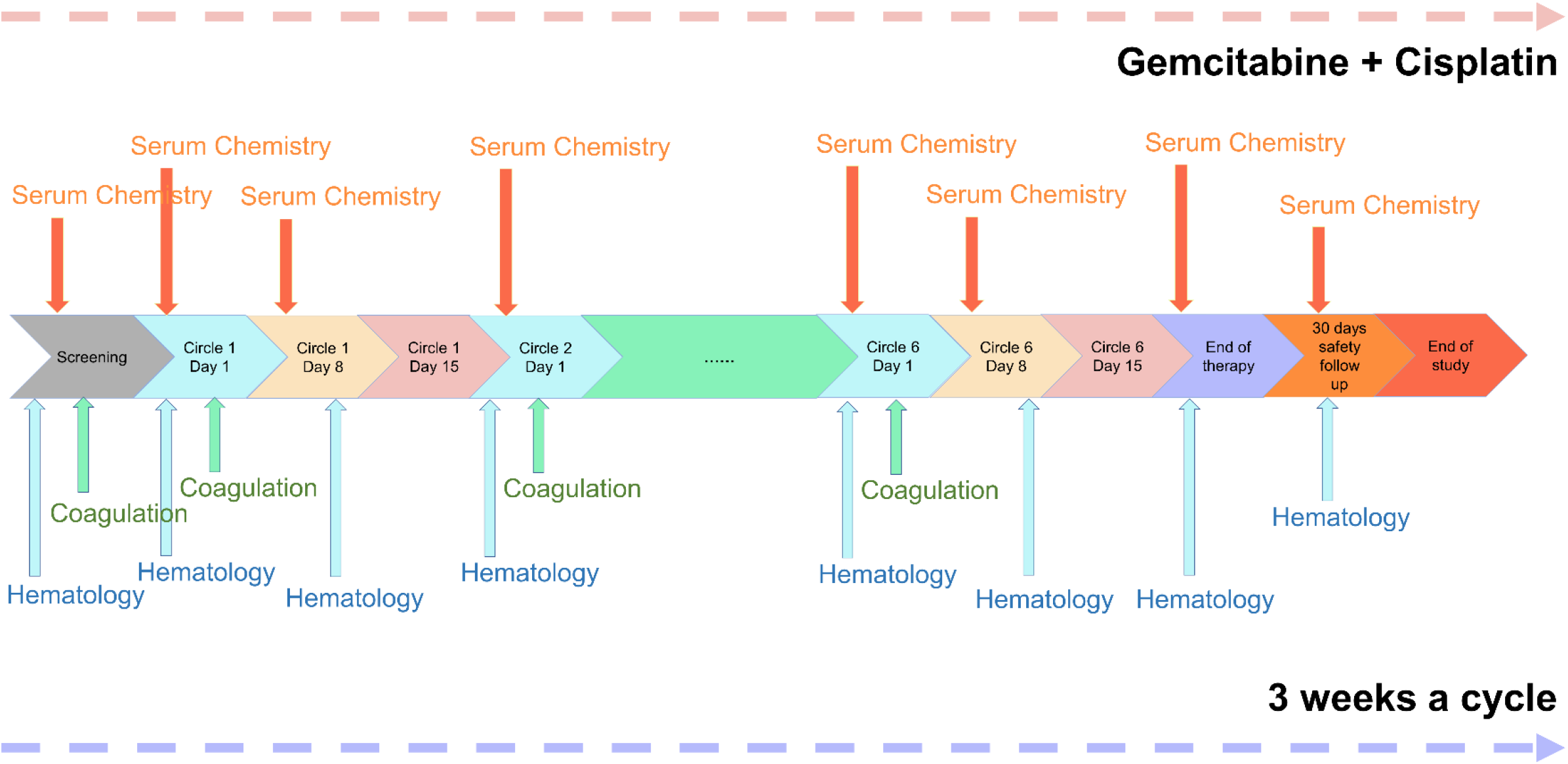
Planned laboratory evaluation.

**Figure S2.**
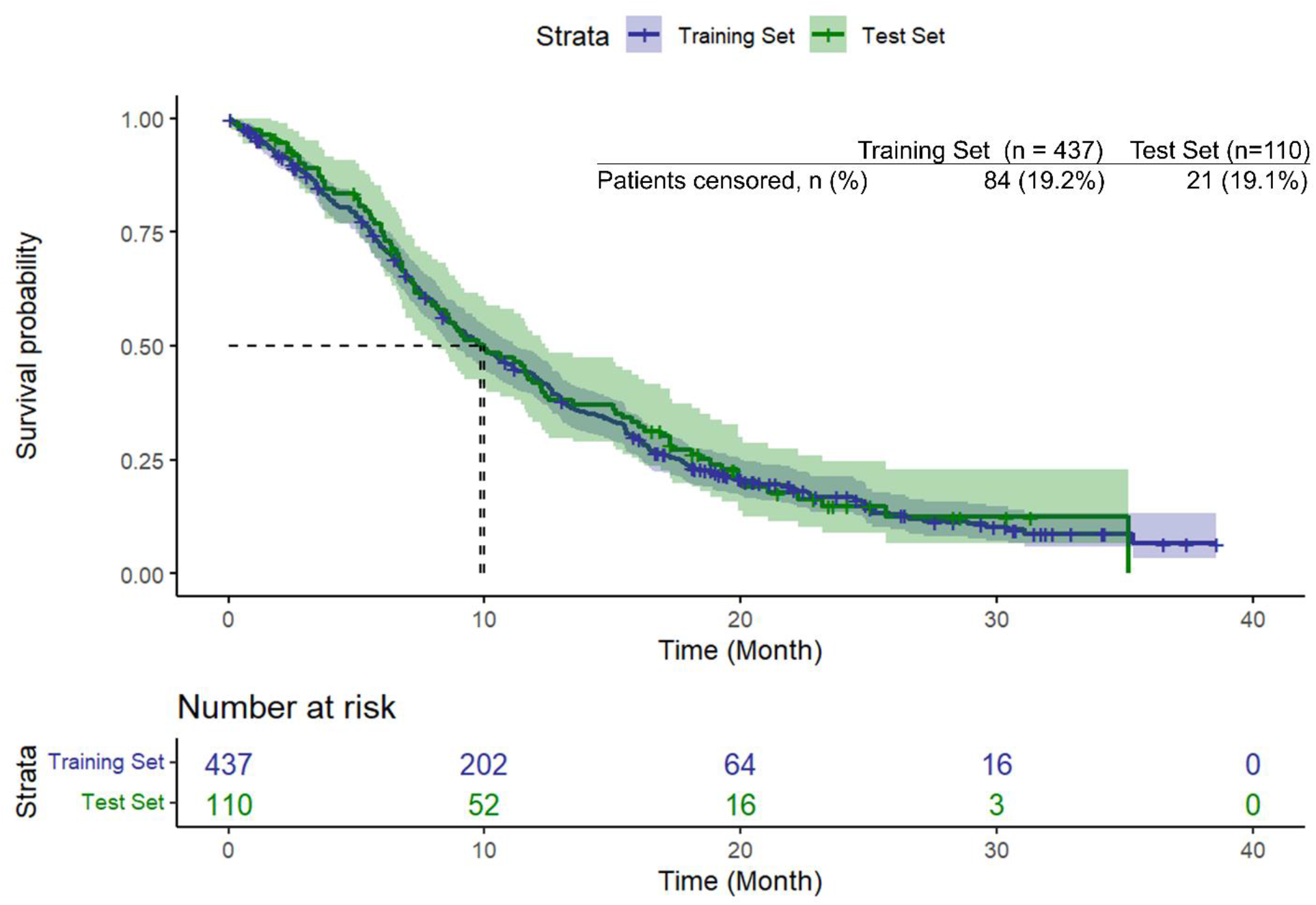
Kaplan-Meier plot for the training and test datasets in a randomly selected cross-validation fold.

**Figure S3.**
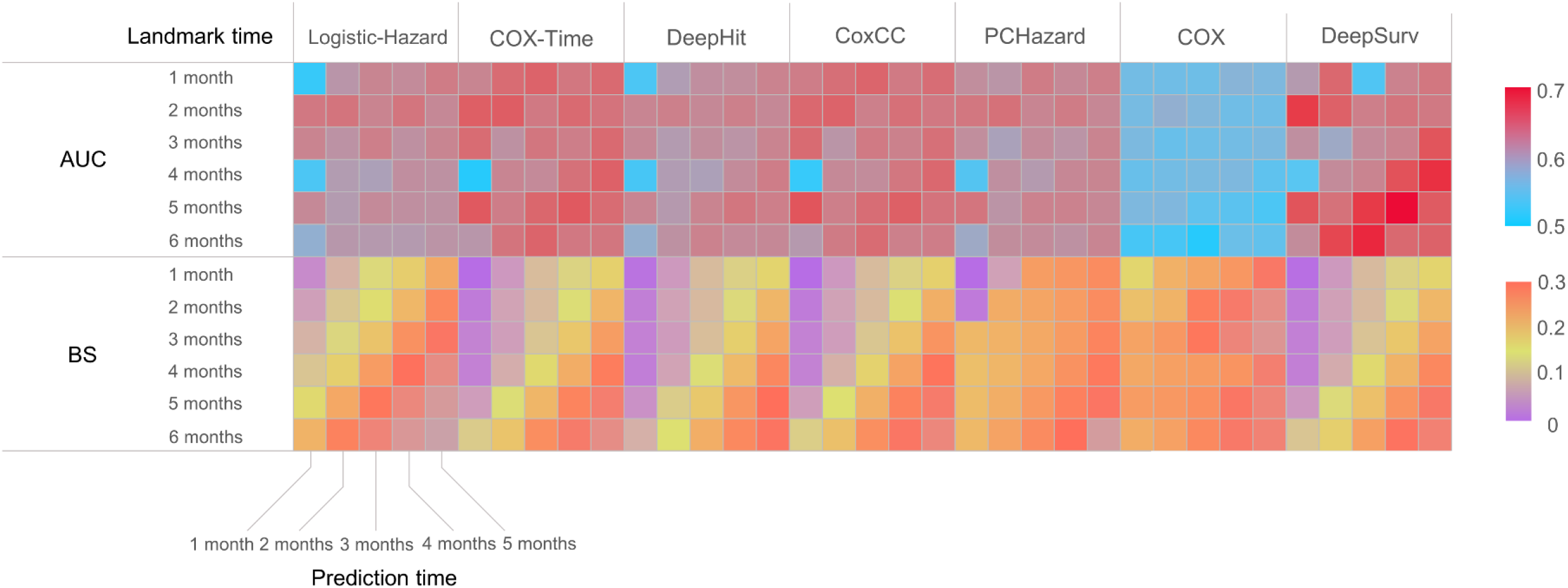
Performance of deep-learning survival models using baseline variables.

**Figure S4.**
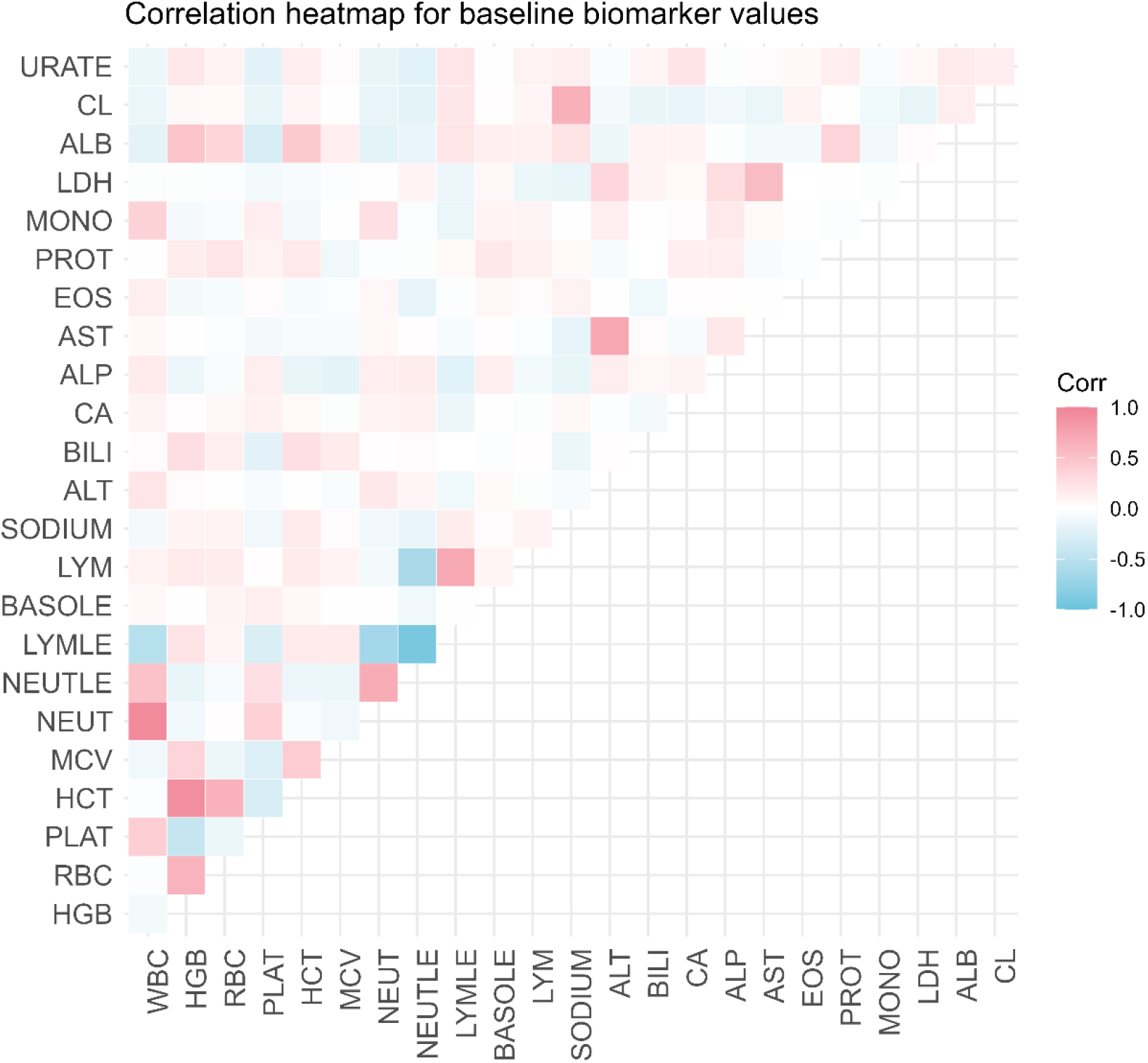
Correlation heat map for baseline biomarker values.

**Figure S5.**
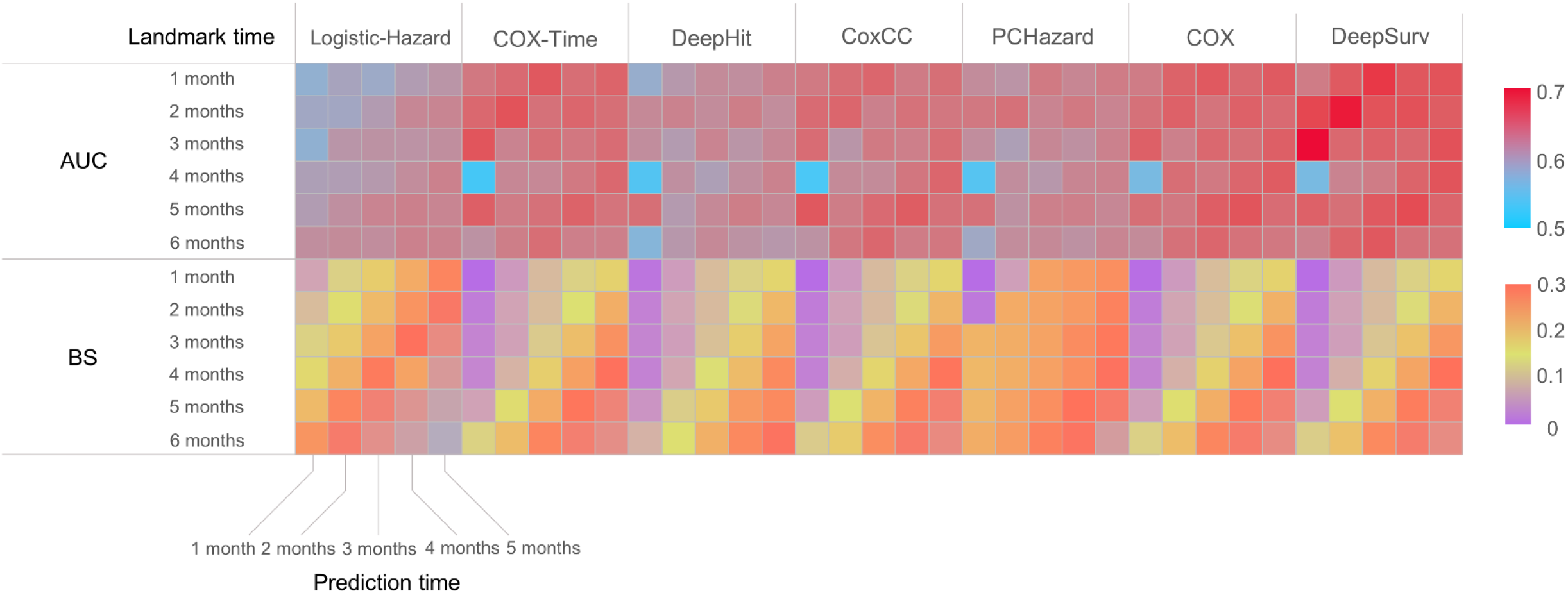
Performance of deep-learning survival models using baseline variables after PCA.

**Figure S6.**
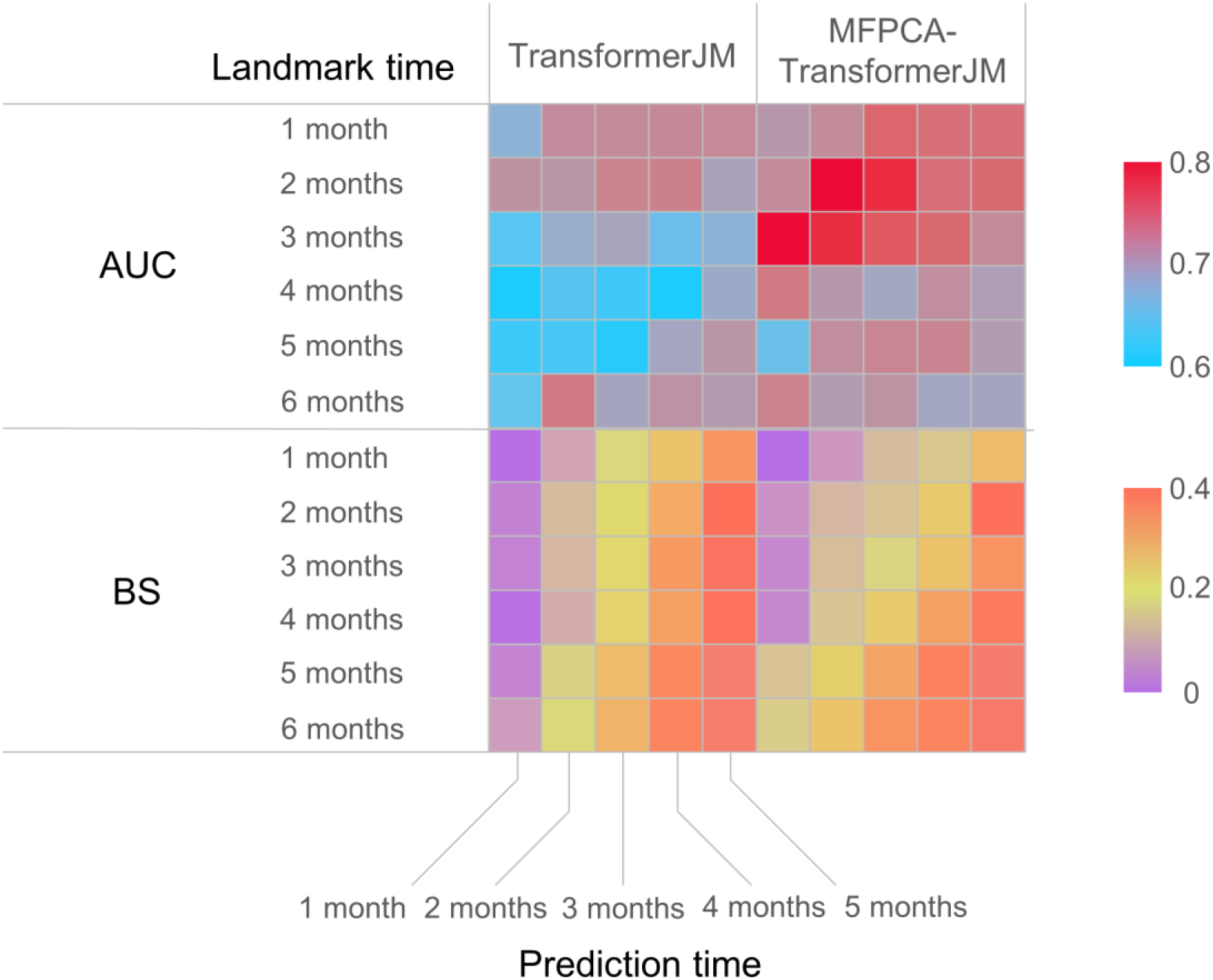
Performance of TransformerJM before and after imputation of longitudinal data using MFPCA.

